# INTRINSIC MOTOR NEURONE EXCITABILITY IS REDUCED IN SOLEUS AND TIBIALIS ANTERIOR OF OLDER ADULTS

**DOI:** 10.1101/2021.06.23.449552

**Authors:** Lucas B. R. Orssatto, David N. Borg, Anthony J. Blazevich, Raphael L. Sakugawa, Anthony J. Shield, Gabriel S. Trajano

**Author notes:** Corresponding author: Lucas Bet da Rosa Orssatto; School of Exercise and Nutrition Sciences, Faculty of Health, Queensland University of Technology (QUT), Brisbane, Australia.

## Abstract

Age-related deterioration within both motor neurones and monoaminergic systems should theoretically reduce neuromodulation by weakening motor neuronal persistent inward current (PIC) amplitude. However, this assumption remains untested. Surface electromyographic signals were collected using two 32-channel electrode matrices placed on *soleus* and *tibialis anterior* of 25 older adults (70±4years) and 17 young adults (29±5 years) to investigate motor unit discharge behaviours. Participants performed triangular-shaped plantar and dorsiflexion contractions to 20% of maximum torque at a rise-decline rate of 2%/s of each participant’s maximal torque. Pairwise and composite paired-motor unit analyses were adopted to calculate delta frequency (ΔF), which has been used to differentiate between the effects of synaptic excitation and intrinsic motor neuronal properties and is assumed to be proportional to PIC amplitude. *Soleus* and *tibialis anterior* motor units in older adults had lower ΔFs calculated with either the pairwise [−0.99 and −1.46 pps; −35.4 and − 33.5%, respectively] or composite (−1.18 and −2.28 pps; −32.1 and −45.2%, respectively) methods. Their motor units also had lower peak discharge rates (−2.14 and −2.03 pps; −19.7 and −13.9%, respectively) and recruitment thresholds (−1.50 and −2.06% of maximum, respectively) than young adults. These results demonstrate reduced intrinsic motor neurone excitability during low-force contractions in older adults, likely mediated by decreases in the amplitude of persistent inward currents. Our findings might be explained by deterioration in the motor neurones or monoaminergic systems and could contribute to the decline in motor function during ageing; these assumptions should be explicitly tested in future investigations.

## INTRODUCTION

The age-related loss of force production has been comprehensively described in the literature [1–3], with the physiological alterations affecting force production including changes in several pathways within the nervous system [1,4–6]. The motor neurone is an important component of the nervous system affected by ageing as it is responsible for integrating and amplifying excitatory synaptic input into an appropriate motor output [7]. An essential intrinsic property of the motor neurone is its capacity to set up persistent inward currents (PICs), which are depolarising currents generated by voltage-sensitive sodium and calcium channels that increase cell excitability by amplifying and prolonging synaptic input [8,9]. Importantly, increases in the concentration of the monoamines serotonin and noradrenaline facilitate PIC development. Under conditions of high monoaminergic drive, synaptic input can be amplified by at least five-fold, suggesting that this amplification is a critical determinant of the motor neurone’s ability to achieve the discharge rates observed during normal motor behaviour [10–12]. Thus, potential physiological alterations in motor neurone intrinsic properties, or in the monoaminergic input to the motor neurone, might reduce the motor neurone’s ability to discharge at higher rates, thus reducing the ability to produce high muscle forces.

During ageing, several changes are observed in the motor neurones that might potentially reduce PIC amplitude in older adults, including lower discharge rates [13,14], reduced incidence of doublet discharges [15], and an increased afterhyperpolarisation duration [16]. These changes are consistent with the lower motor neurone excitability that is also observed in aged rat models [17,18]. With respect to the monoaminergic system, research using both human and animal models suggests that ageing is associated with reduced noradrenaline and serotonin secretions and thus input onto the motor neurones [19–23], which might theoretically underpin PIC amplitude reduction with ageing. These findings indicate the possibility that PIC amplitude might be reduced in older adults; however, this hypothesis remains to be tested [24].

The amplitude of PICs can be estimated in humans using the paired motor unit technique [8,25,26], with data obtained using high-density surface electromyography [27,28]. This technique requires the pairing of the discharge rates of a low-threshold (control unit) to a higher-threshold (test unit) motor unit, obtained during a slowly-increasing and decreasing triangular-shaped contraction [8,25,29]. Subsequently, the difference in discharge rate of the control unit between the time of recruitment and de-recruitment of the test unit is computed as the change in frequency (ΔF). ΔF has been used to differentiate between the effects of synaptic excitation and motor neuronal intrinsic properties and is assumed to be proportional to PIC amplitude [9]. However, ΔF values need to be interpreted with caution as they can be affected by spike frequency adaptation, spike frequency accommodation, and the proportion of sub-threshold PICs [8,30,31]. When controlling for these confounding factors, the technique can be used to estimate and compare PIC amplitudes in motor units of young and older adults.

The present study compared ΔF amplitudes (i.e., estimates of PICs) of *soleus* and *tibialis anterior* motor units between young and older adults. Additionally, we explored the relationship between ΔF and the peak discharge rates. We hypothesised that there would be a reduction in older adults’ ΔF in both *soleus* and *tibialis anterior,* and that ΔF would be strongly associated with motor unit peak discharge rate. *Soleus* and *tibialis anterior* were selected for study because the control and timing of their activation are both critical to the performance of daily activities such as standing and walking in older adults [32,33].

## METHODS

### Participants and ethical procedures

Forty-four participants were recruited for the study, including 18 young adults and 26 older adults (participant characteristics are documented in Table 1). More older adults were recruited because it was expected that fewer motor units would be identified during decomposition and that some participants may not be able to perform the triangle-shaped contractions with the necessary torque rise and fall accuracy. To participate, volunteers had to: a) be young adults aged 18 – 35 years or older adults ≥65 years; b) have no history of neurological disorders; c) be free of lower limb musculoskeletal injuries; and d) not be taking medications that could influence the monoaminergic system, including serotonin or noradrenaline modulators (e.g., beta-blockers and serotonin reuptake inhibitors). Also, participants were instructed to not consume caffeinated foods (e.g., coffee) 24 h before the testing session. Participants were excluded from the analyses if: a) no usable motor units were identified by the decomposition algorithm, or b) if it was not possible to achieve all the assumptions required in the paired motor unit analysis (as described below), for at least one pair of motor units, for either *soleus* or *tibialis anterior.* One participant per group was excluded from the study because no motor units were identified in either *soleus* or *tibialis anterior.* The study was approved by the University Human Research Ethics Committee, and all participants gave written informed consent before participating. Data collection was conducted during the COVID-19 pandemic and all safety procedures followed the local state government policies.

**Table 1.** Participant characteristcs.

| Variable | Young Adults (n = 17) | Older Adults (n = 25) |
| --- | --- | --- |
| Age (years) | 29 ± 5 | 70 ± 4 |
| Sex (n, %) |  |  |
| Men | 9 (53%) | 11 (44%) |
| Women | 8 (47%) | 14 (56%) |
| Body mass (kg) | 71.9 ± 13.6 | 77.8 ± 19.3 |
| Body fat (%) | 20.2 ± 7.7 | 33.0 ± 10.0 |
| Appendicular skeletal muscle mass (kg) | 25.2 ± 6.7 | 21.6 ± 5.1 |
| Height (cm) | 175 ± 11 | 166 ± 9 |
| Handgrip strength (kg) | 40.3 ± 13.3 | 26.4 ± 10.5 |
| Peak torque (N·m) |  |  |
| Plantar flexion | 156 ± 47 | 85 ± 32 |
| Dorsiflexion | 41 ± 14 | 29 ± 7 |
| Normalised peak torque (N·m·kg<sup>-1</sup>) |  |  |
| Plantar flexion | 2.13 ± 0.49 | 1.11 ± 0.37 |
| Dorsiflexion | 0.56 ± 0.09 | 0.38 ± 0.10 |
| Functional capacity |  |  |
| Timed up-and-go (s) | - | 6.1 ± 1.0 |
| 5-times sit-to-stand (s) | - | 13.4 ± 2.9 |
| Physical activity level (MET-min/week) | 2541 (1902–3039) | 2994 (1386–5139) |

Note: Data are presented as mean ± standard deviation, except for Physical activity level, which is presented as median with interquartile range. MET-min/week, metabolic equivalent of task score of the performed physical activities by the minutes accomplished per week.

### Study design and testing procedures

Participants visited the laboratory on a single occasion in which they were familiarised with the testing procedures and data were collected. Initially, participants signed the informed consent form, and completed the International Physical Activity Questionnaire (short-version), which was used to estimate weekly physical activity levels based on the metabolic equivalent of a task (MET – i.e., multiples of the resting metabolic rate) for each physical activity domain. MET scores for each activity were multiplied by the minutes performed in one week, providing the total MET-min/week [34,35]. Physical activity levels were interpreted based on the recommendations of the Guidelines for Data Processing and Analysis of the International Physical Activity Questionnaire (IPAQ) [35]. Thereafter, body composition assessment was conducted with a multi-frequency bioelectrical impedance device (MC-780, Tanita, Japan), which provided body fat percentage and appendicular skeletal muscle mass data.

After electrode placement on *soleus* and *tibialis anterior,* the participants were seated upright in the chair of an isokinetic dynamometer (Biodex System 4, Biodex Medical system, Shirley, NY) with the knee fully extended (0°) and ankle in the anatomical position (0°). A warm-up consisting of six 5-s submaximal voluntary isometric plantar and dorsiflexion contractions (2 × 30%, 2 × 60%, and 2 × 80% of perceived maximal effort) was performed, followed by three maximal voluntary contractions of ~3-s with 30-s rest intervals. The maximum torque achieved was recorded as the maximal voluntary contraction peak torque, which was also normalised to body mass. Subsequently, participants were familiarised with the triangular-shaped contractions to 20% of their maximum voluntary torque level. Triangular contractions to 20% of maximal torque have been extensively used for ΔF calculations using the paired motor unit technique [8,31,36–38], and this force was considered similar to the average torques developed during daily activities such as standing [39] and walking. All contractions had a duration of 20 s (10-s up and 10-s down) and were performed at a rate of torque increase and decrease of ~2%/s. Participants were instructed to follow the torque path provided in real time on a 58-cm computer monitor during each contraction. Data collection commenced 5 min after the end of familiarisation (usually requiring ~3-10 × 20% triangular contractions with 30-s rest), during which the participants then performed four triangular contractions with 60-s rest intervals. When an abrupt increase or decrease in torque was observed (i.e., the torque trajectory was not closely followed), the trial was excluded and repeated. The maximum voluntary isometric torque and order of triangular contraction completion was randomised between *soleus* and *tibialis anterior.*

After the neuromuscular assessments, the participants performed a handgrip strength test using a grip force transducer (ADinstruments, Australia). They performed 3 submaximal (50% of their perceived maximal effort) familiarisation contractions, followed by 3 ×3-s maximal contractions with 30-s rest intervals. The maximum force achieved was recorded as their handgrip strength. Thereafter, they performed sit-to-stand and timed up-and-go (functional capacity) tests, timed off-line with video recordings as recommended by da Silva et al. [40] to reduce measurement error. The 5-times sit-to-stand required the participants to stand up five times until they reached upright standing and then returned to the seated posture on a chair (seat 46 cm high). The timed up-and-go test required the participants to rise from a chair (seat 46 cm high), walk towards and around a cone 3 m from the chair, return to the starting position, and to sit without the aid of hands and not running, in the shortest possible time. The fastest of three attempts (60-s rest between) was analysed.

The participants’ sarcopaenia status was screened according to the cut-off points and algorithm from the European Working Group on Sarcopenia in Older People (EWGSOP2) [48]. According to this scale, older adults with low handgrip strength (<27 kg for men or <16 kg for women) are classified as “sarcopaenia probable”, and if this is accompanied by low appendicular skeletal muscle mass (<20 kg for men and <15 kg for women) then they are confirmed as sarcopaenic. Older adults with confirmed sarcopenia who take longer than 20 s to perform the timed up-and-go test are classified as having severe sarcopaenia. Also, participants with timed up-and-go performance under 12 s are classified as having normal mobility [41].

### Surface electromyography

Surface electromyograms (sEMG) were recorded during the 20% triangular contractions using four semi-disposable 32-channel electrode grids with a 10-mm interelectrode distance (ELSCH032NM6, OTBioelettronica, Torino, Italy). After skin shaving, abrasion, and cleansing with 70% isopropyl alcohol, two electrode grids were placed over the medial and lateral portions of *soleus* (either side of the Achilles tendon) and another two electrode grids were placed over the superior and inferior aspect of *tibialis anterior* using a bi-adhesive foam layer and conductive paste (Ten20, Weaver and Company, Colorado, USA). A strap electrode (WS2, OTBioelettronica, Torino, Italy) was dampened and positioned around the ankle joint as a ground electrode. The sEMG signals were acquired in monopolar mode, amplified (256×), band-pass filtered (10–500 Hz), and converted to a digital signal at 2048 Hz by a 16-bit wireless amplifier (Sessantaquattro, OTBioelettronica, Torino, Italy) using OTBioLab+ software (version 1.3.0., OTBioelettronica, Torino, Italy) before being stored for offline analysis.

### Motor unit analyses

#### Motor unit identification

The recorded data were processed offline using the DEMUSE software [27]. For each muscle, only the triangular contraction yielding the lowest deviation from the torque trajectory was analysed. If both contractions presented a similar torque trajectory, the contraction with the highest number of identified motor units was analysed. High-density sEMG signals were band-pass filtered (20-500 Hz) with a second-order, zero-lag Butterworth filter. Thereafter, a blind source separation method, the convolutive kernel compensation (CKC) method, was used for signal decomposition [27,42] from each triangular contraction. CKC yields the filters of individual motor units (so-called motor unit filters) that, when applied to high-density sEMG signals, estimate the motor unit spike trains [27,42]. After decomposition, a trained investigator manually inspected motor unit spike trains and edited the discharge patterns of the motor units. Only the motor units with a pulse-to-noise ratio equal to or greater than 30 dB were kept for further analysis [42].

#### Estimation of PIC amplitude (ΔF) and peak discharge rate

The observed discharge events for each motor unit were converted into instantaneous discharge rates and fitted into a 5^th^-order polynomial function. The maximum value obtained from the polynomial curve was considered the peak discharge rate. The relative torque (%) produced at the time in each motor unit was recruited was considered the recruitment threshold. The recruitment threshold was used to characterise the populations of motor units identified by the decomposition algorithm for each group. Thereafter, PIC amplitude was estimated using the paired motor unit analysis [8], referred through the manuscript as pairwise method. Motor units with a low recruitment threshold (i.e., control units) were paired with higher recruitment threshold motor units (i.e., test units). ΔF was calculated as the change in discharge rates of the control motor unit from the moment of recruitment to the moment of de-recruitment of the test unit [8,9]. In order to produce motor unit pairs, the following criteria were adopted: 1) rate-to-rate correlations between the smoothed discharge rate polynomials of the test and control units was *r* ≥0.7; 2) test units were recruited at least 1.0 s after the control units; and 3) the control unit did not show discharge rate saturation after the moment of test unit recruitment (i.e., discharge rate from the control unit at the moment the test unit was recruited minus the peak discharge rate at the control unit >0.5 pps) [8,38,43–45]. ΔFs obtained for each control unit were averaged to obtain a single ΔF for each test motor unit.

We also conducted an additional analysis using the composite paired motor unit method to calculate ΔF values for each motor unit [31]. This method is characterised by the overlay of 3 lower-threshold motor units to construct a single composite control unit profile to be paired with the test units. The composite method has been suggested to address some of the limitations observed with the pairwise method, such as underestimation and overestimation of ΔF values, reducing its variability. However, strict assumptions are made for eligible motor units to be included in the analysis, which allows its use only in muscles in which it is possible to identify many motor units (e.g., *tibialis anterior)* but not in those in which fewer motor units are identified using the decomposition method (e.g., *soleus’).* Moreover, this method does not allow calculation of ΔF values for lower-threshold motor units since they are used to construct the composite control unit. In summary, the control unit is the overlay of the instantaneous discharge rate of at least three motor units recruited at <3% of the maximum voluntary torque and presenting a similar discharge profile, which was determined visually by an experienced researcher with individual motor units discharge rates plotted superimposed to each other. This is followed by the application of the 5^th^ order polynomial over the overlayed motor units discharge rates. It was assumed that lower-threshold motor units had their discharge rate profile more linearly related to the synaptic input profile because PICs were almost fully activated at the time of recruitment; therefore, it would be more appropriate to use the composite unit as control. Also, this method requires the removal of the acceleration phase of the discharge rates (i.e., secondary range), making the polynomial ascending-to-descending slope ratio near 1, which is important when measuring PIC amplitudes to avoid any rate-dependent effects on motor unit recruitment or derecruitment [46]. The secondary range was determined with visual inspection of deflection point following Afsharipour et al., [31] procedures. The test units should start discharging in the tertiary range (i.e., after the secondary range and before the descending phase). This method provides a single ΔF for each motor unit, which was then used in the data analysis. Figure 1 illustrates the pairwise and composited paired motor unit analysis methods on *tibialis anterior* motor units for one participant per group. Panels C and D display the test units used for both methods. Panels E and F display the control units for the pairwise method and panels G and H display the control units for the composite method.

**Figure 1.**
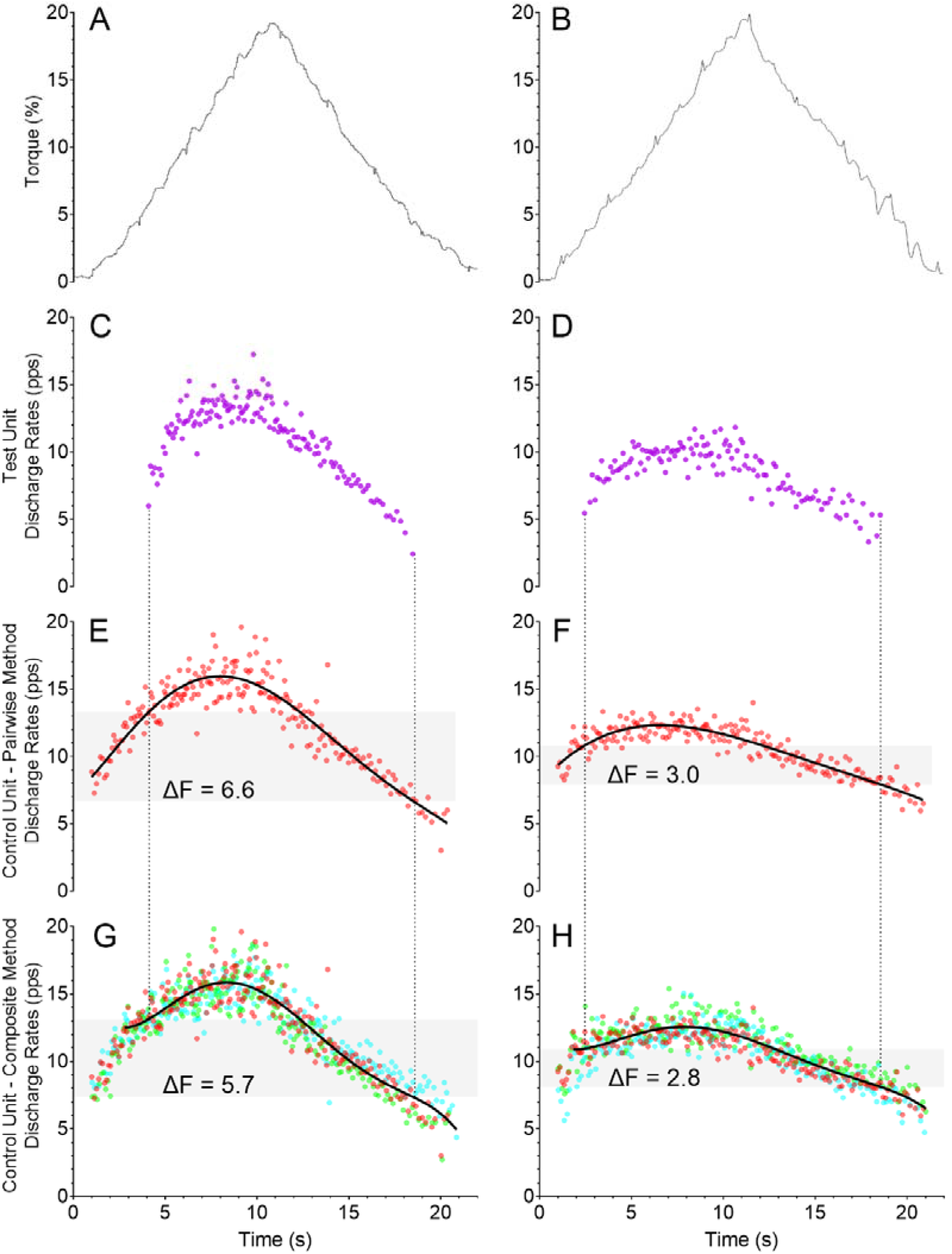
Data from a single participant for each group showing torque during triangle-shaped contractions and delta frequency (ΔF) calculation in *tibialis anterior* for both the pairwise and composite paired motor unit analyses. Data from a young adult is displayed in the left panels and from an older adult in the right panels. Panels A and B on the first row show the torque traces for contractions with 20% of the participant’s maximal voluntary torque. The participants’ test units are displayed on panels C and D (purple motor unit), control units for the pairwise method on panels E and F (red motor unit), and control units for the composite method on panels G and H (red, green, and blue motor units). The black continuous lines are the 5^th^-order polynomial fits for the control units. Note that for the composite method the polynomial curve starts from the tertiary range. The gray-shaded areas represent the ΔF amplitude for each participant and analysis.

### Data analysis

All analyses were undertaken in R (version 4.0.3) using RStudio environment (version 1.3.1093). Models were fitted using the *lmerTest* package [47]. A linear mixed-effects model was used to compare estimates of ΔF of *soleus* and *tibialis anterior* motor units between young and older adults. The model included: age group, muscle type and recruitment threshold as fixed effects. A random intercept and correlated random slopes (recruitment threshold and muscle type) were included for each participant in the study, to account for the correlation between repeated observations on each individual. This model was selected from a series of candidate models (Supplement 1), based on the smallest Bayesian Information Criteria value. The recruitment threshold was standardised (mean = 0, SD = 1) before analysis.

Separate linear mixed-effects models were used to analyse peak discharge rate and recruitment threshold data. These models included: age group, muscle type and age group by muscle type as fixed factors; and a random intercept and slope (muscle group) for each participant. The estimated marginal mean difference and 90% and 95% confidence intervals (CI) in ΔF, peak discharge rate, and recruitment threshold between young and older adults, were determined using the *emmeans* package [48]. The standardised difference, denoted *d*, was also calculated using the population SD from each respective linear mixed-effects model as the denominator [48].

To determine the contribution of ΔF to peak discharge rate, a linear mixed-effects model was fitted, with the coefficient of determination (R^2^) used to quantify the proportion of the variance in peak discharge rate explained by ΔF [49]. The model included ΔF as a fixed effect and a random intercept and slope (ΔF) for each participant, to account for the correlation of repeated measurements on an individual. ΔF was standardised (mean = 0, SD = 1). Differences between young and older adults in peak plantar flexion and dorsiflexion torque, and physical activity levels, were determined using independent *t*-tests. The α level for all tests was 5%. The dataset and R code can be found at https://github.com/orssatto/PICs-ageing.

## RESULTS

### Effects of age and muscle group on ΔF, peak discharge rate, and recruitment threshold

Motor units of older adults identified by the decomposition algorithm in our study had lower ΔFs and peak discharge rates and were recruited at lower torque (muscle force) levels than young adults in both *soleus* and *tibialis anterior* (Figure 2, panels A, B, and C, respectively). Also, ΔF levels and peak discharge rates were lower in *soleus* than *tibialis anterior,* independent of age (Figure 2, panels A and B).

**Figure 2.**
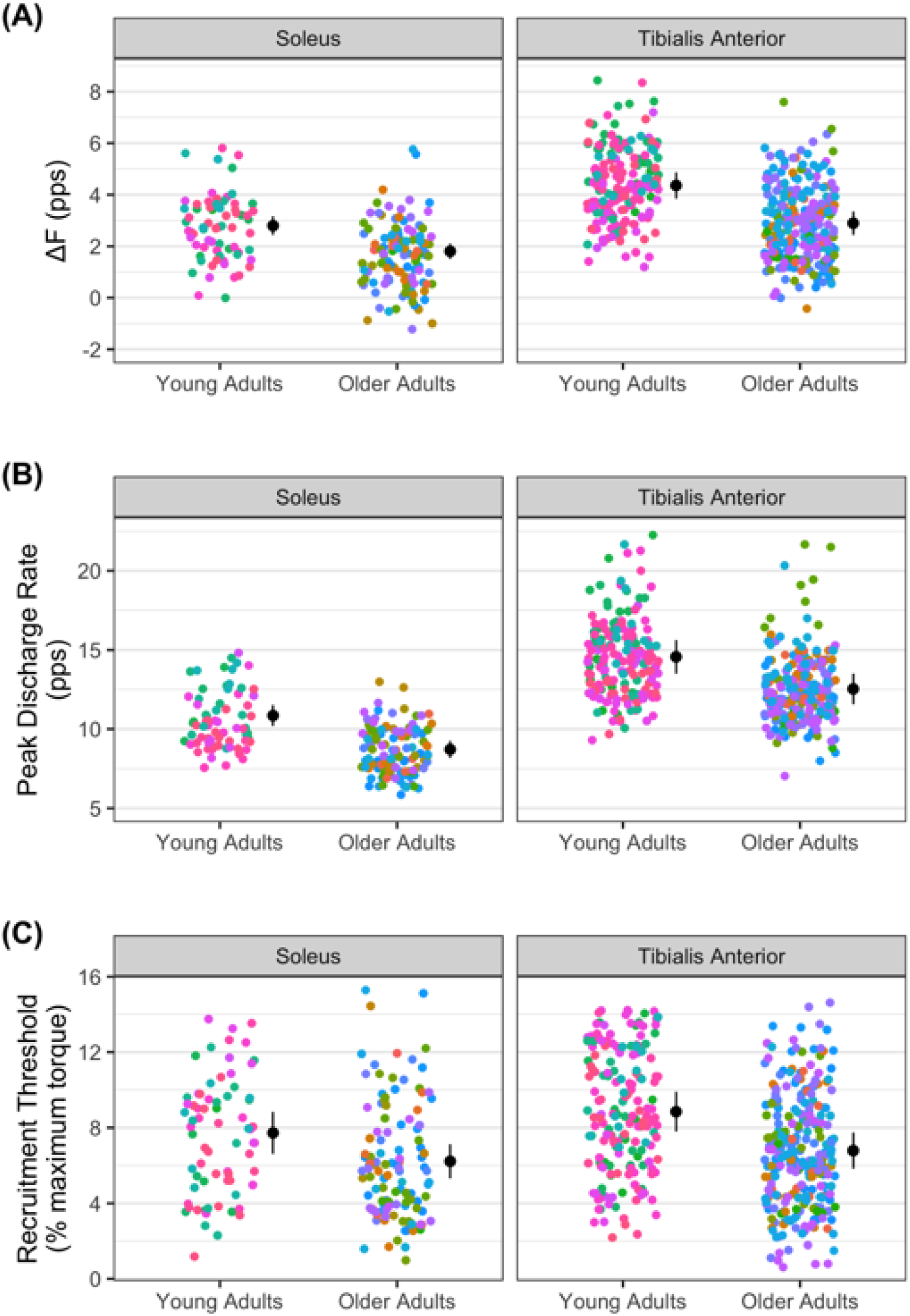
ΔF calculated with the pairwise paired motor unit method (A), peak discharge rate (B), and recruitment threshold (C) in *soleus* and *tibialis anterior* in young and older adults. The mean (circle) and 95% confidence interval are offset to the right, with individual data points coloured by participant. pps = peaks per second.

There were effects of age *(β* = −0.99, 95% CI = −1.42, −0.57;*p* < .001), muscle *(β* = 1.56, 95% CI = 1.05, 2.07; *p* < .001) and recruitment threshold (*β* = 0.47, 95% CI = 0.27, 0.66; *p* < .001) but no age group by muscle effect *(β* = −0.47, 95% CI = −1.15, 0.21; *p* = .18), on ΔF when calculated using the pairwise paired motor unit method. ΔF was lower in older adults (Figure 2A) in both *soleus (d* = −0.93; Figure 3A) and *tibialis anterior* (*d* = −1.38; Figure 3A). There were effects of age (*β* = −2.14, 95% CI = −2.98, −1.33; *p* < .001) and muscle (*β* = 3.71, 95% CI = 2.64, 4.83; *p* < .001) on peak discharge rate, but there was no age by muscle effect on peak discharge rate (*β* = 0.11, 95% CI = −1.37, 1.56; *p* = .88). Peak discharge rate was lower in older adults (Figure 2B) in both *soleus* (*d* = −0.98; Figure 3B) and *tibialis anterior (d* = −1.27; Figure 3B). There was an age effect on recruitment threshold, with thresholds lower in older adults *(β* = −1.50, 95% CI = −2.89, −0.12; *p* = .040; Figure 3C). There was no evidence of muscle *(β* = 1.12, 95% CI = −0.35, 2.65; *p* = .15) or age by muscle *(β* = 1.12, 95% CI = −2.59, 1.38; *p* = .58) effects on recruitment threshold.

**Figure 3.**
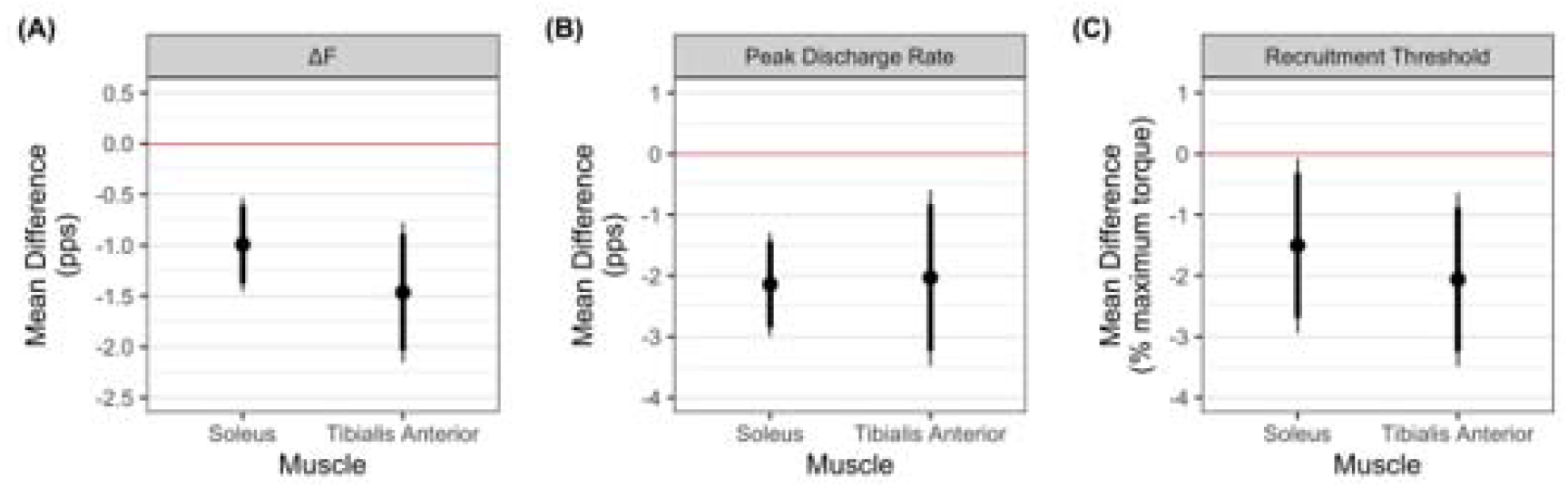
The marginal mean difference with 90% (thick line) and 95% (thick line) confidence intervals between young and older adults for ΔF calculated with the pairwise paired motor unit method (A), peak discharge rate (B), and recruitment threshold (RT) (C) in *soleus* and *tibialis anterior.* Negative values on all panels indicate the lower responses in older adults. There was an effect of age on ΔF, peak discharge rate, and recruitment threshold, with all variables lower in older adults. There was no age by muscle effect on ΔF, peak discharge rate, or recruitment threshold—indicating that age-dependent differences in these variables were similar for both muscles.

The results of the additional, composite method, analysis revealed there was effects of age *β* = −1.18, 95% CI = −1.85, −0.51;*p* = .002), muscle *β* = 1.37, 95% CI = 0.74, 2.00;*p* < .001), and age by muscle *(β* = −1.10, 95% CI = −1.39, −0.27; *p* = .014) on ΔF. ΔFs were lower in older adults in both *soleus (d* = −1.54) and *tibialis anterior (d* = −2.98), as shown in Figure 4. Also, ΔFs were lower in *soleus* than *tibialis anterior* in young adults (mean difference = −1.37, 95% CI = −2.27, −0.46; *p* = .008; *d* = −1.79) but not older adults (mean difference = −0.27, 95% CI = −1.04, 0.51; *p* = .47; *d* = −0.35). The composite method removed the effect of recruitment threshold on ΔF *(β* = −0.05, *p* = .61) that was observed when modelling the pairwise method data *(β* = 0.47, *p* < .001).

**Figure 4.**
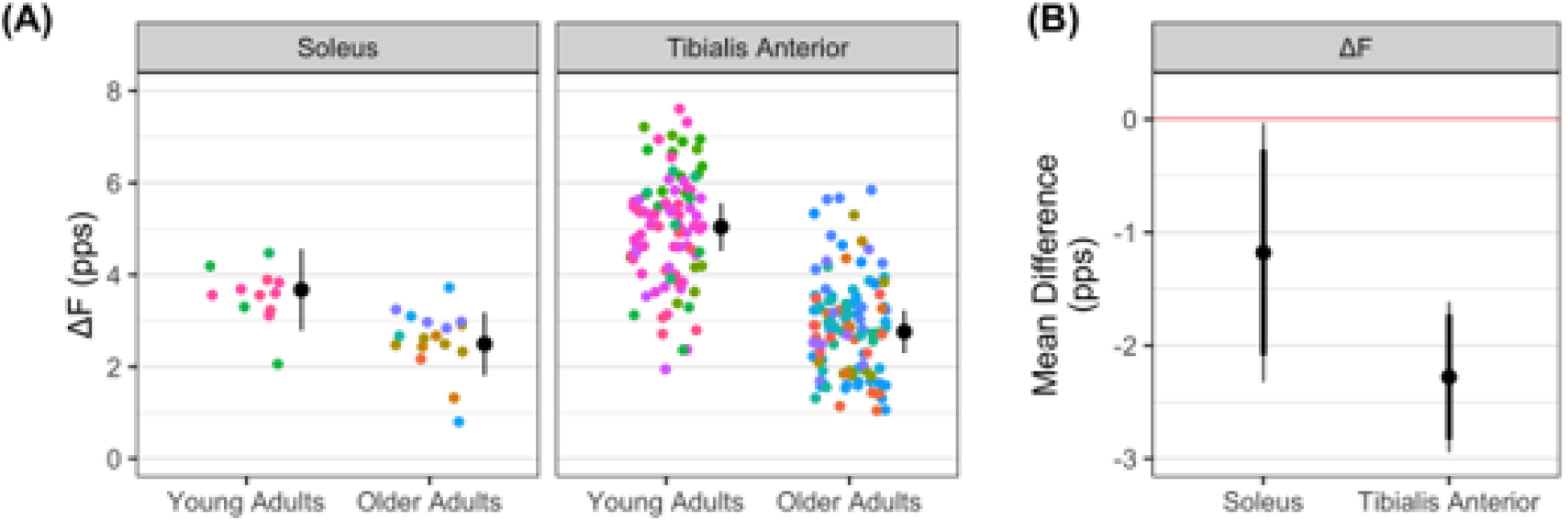
ΔF calculated with the composite paired motor unit method (A), and the marginal mean difference with 90% (thick line) and 95% (thick line) confidence intervals between young and older adults for ΔF (B). In panel A, the mean (circle) and 95% confidence interval are offset to the right, with individual data points coloured by participant. In panel B, negative values indicate ΔF was lower responses in older adults. pps = peaks per second.

The analysis of the contribution between ΔF and peak discharge rate showed that, when accounting for the correlation between repeated observations on each participant, ΔF explained 53% of the variance in peak discharge rate. There was a statistical effect of ΔF on peak discharge rate *(β* = 1.19, 95% CI = 0.87, 1.51, *p* = .001).

### Participant characteristics

Participant characteristics are shown in Table 1. Body fat (%) was higher (mean difference = 12.7% of body mass, 95% CI = 7.1, 18.4; p < .001; *d* = 1.40) and appendicular skeletal muscle mass was lower (mean difference = −3.7 kg, 95% CI = −7.3, −0.1; *p* = .045; *d* = −0.63) in older adults than young adults. Handgrip strength was lower in older adults (mean difference = −14.0 kg, 95% CI = −21.2, −6.7; *p* < .001; *d* = −1.19). Absolute peak torque was lower in older adults in both plantar flexion (mean difference = −70 N·m, 95% CI = −95, −46; *p* < .001; *d* = −1.83) and dorsiflexion (mean difference = −12 N·m, 95% CI = −18, −5; *p* < .001; *d* = −1.18), as was the normalised peak torque in both plantar flexion (mean difference = −1.02 N·m·kg^-1^, 95% CI = −1.28, −0.77; *p* < .001; *d* = −2.45) and dorsiflexion (mean difference = −0.17 N·m·kg^-1^, 95% CI = −0.23, −11; *p* < .001; *d* = −1.81).

According to the EWGSOP2 sarcopaenia cut-off points and algorithm [50], no participant was confirmed as sarcopaenic. Three older men and 1 older woman were classified as “sarcopaenia probable” (handgrip strength <27 kg for men and <16 kg for women), but their appendicular skeletal muscle mass levels were higher than the cut-off points, not confirming sarcopaenia (>20 kg for men or >15 kg for women). All older adults performed the timed up-and-go test in < 12 s and were classified as normal mobility.

Physical activity levels were not statistically different between young and older adults (mean difference = 594 MET min/week, 95% CI = −1269, 2457; *p* = .52; *d* = 0.20), with both young and older adults presenting with moderate-to-high physical activity levels.

### Motor unit identification

The number of motor units identified in *soleus* was 113 for the young group and 211for the older group. The median (IQR) number per younger participant was 7 (5-8) and per older participant 8 (6-9). For *tibialis anterior,* 293 motor units were identified in the young group and 411 in the older group. The median number per younger participant was 19 (16-23) and per older participant 20 (16-27). For the pairwise paired motor unit analysis, it was possible to obtain ΔF values from 16 young adults and 23 older adults in *soleus,* and from 16 young adults and 19 older adults in *tibialis anterior.* The number of test unit ΔFs for *soleus* in the young group was 70 and in the older group was 117. The median number per young participant was 5 (3-6) and per older participant was 4 (3-6). The number of test unit ΔFs for *tibialis anterior* in the young group was 185 and in the older group was 257. The median number per younger participant was 11 (8-14) and per older participant was 14 (9-18).

For the composite paired motor unit analysis, it was possible to obtain ΔF values from 4 young adults and 7 older adults for *soleus* and from 15 young adults and 18 older adults for *tibialis anterior.* The number of test unit ΔFs for *soleus* in the young group was 12 and in the older group was 17, with the median number per younger participant being 3 (3–4) and per older participant being 2 (1–4). The number of test unit ΔFs for *tibialis anterior* in the young group was 93 and in the older group was 111, with the median number per young participant being 5 (4–8) and per older participant being 6 (3–8). No motor units were identified in one young participant and one older participant in *soleus,* and in one young participant and six older participants in *tibialis anterior,* respectively. These participants were not included in the analyses for the respective muscles.

## DISCUSSION

The present study compared amplitudes of motor neuronal PICs, estimated using the paired motor unit technique (ΔF), between young and older adults. The main findings indicate that ΔF values are considerably lower in both *soleus* and *tibialis anterior* in older adults. These reductions are accompanied by lower peak discharge rates and the recruitment of motor units at lower torque levels in the older than younger adults in both muscles. As an exploratory analysis, a model was developed that showed a small contribution of ΔF to the between-subject variance in peak discharge rates. These findings point to a meaningful reduction in the intrinsic motor neurone excitability in a group of non-sarcopaenic older adults with normal mobility that is at least partly associated with a reduced amplitude of persistent inward currents.

### Estimates of persistent inward currents (ΔF)

The ΔF values obtained from *soleus* and *tibialis anterior* in young adults are similar to those obtained in previous studies using triangular-shaped contractions with a 2%/s force increase-decrease rate [31,36,37] and indicate that ΔFs from older adults are substantially lower than in young adults in both *soleus* and *tibialis anterior.* However, our results partially differ from previous findings that investigated self-sustained discharging in older humans [51] and the frequency-current slope in aged rats [18]. Kamen et al., [51] observed no difference between young and older adults in self-sustained discharging of *tibialis anterior* motor units when excitatory input was applied by delivering a brief vibration stimulus. In addition, they have observed no differences between age groups in maximal voluntary force and no change in force caused by the vibration stimulus. These results could indicate that the reductions in PICs with ageing seem clearer when tested with the paired-motor-unit analysis, or because older adults present significantly lower force levels than their younger counterparts. In agreement to our result, Kalmar et al., [18] observed lower frequency-current slope, which indicate lower PICs amplitude, in older rats hindlimb muscles. However, they have also reported higher incidence of PICs in older rats, which could be an adaptative response to counteract the reductions in PICs amplitude, justified by augmentation of the serotonin and noradrenaline receptor sensitive to residual endogenous monoamines [52].

Weaker PICs in older adults can be a result of the detrimental changes within the monoaminergic systems, which indicate lower release of serotonin and noradrenaline with ageing. Locus coeruleus and the dorsal raphe nucleus are the major central sources of noradrenaline and serotonin, respectively [53]. Diminution of structural integrity [19,54] and in neuromelanin (a product of noradrenaline synthesis) content in noradrenergic neurones emanating from locus coeruleus [20] indicate impaired noradrenaline secretion in older adults. Moreover, noradrenaline and serotonin concentration decrements have been observed in the brains of aged rats [21,22], and degeneration of serotonergic axons projecting to the ventral horn of the lumbar segment of the spinal cord (where motor neurones innervating lower limb muscles emanate) have been detected [23]. Also, serotonin receptors are affected by expanded circulation of cytokines, resulting in increased re-uptake of serotonin [55]. Thus, older adults might speculatively present reduced noradrenaline and serotonin secretions, and hence input onto motor neurones, which would then impair the initiation and modulation of PICs in this population. Changes in motor neurone integrity might also partly underpin reductions in intrinsic excitability. During ageing, axonal demyelination due to reduced expression of proteins responsible for myelination [56] as well as axonal atrophy and degeneration have been observed, possibly subsequent to deregulated Ca^2+^ homeostasis [57] and to toxic, metabolic, or infectious injury sustained throughout the lifespan, or due to high levels of chronic inflammation and oxidative stress [58,59]. Motor neuronal death, especially in higher-threshold motor axons, leading to denervation of motor units has also been documented [60]. In these cases, denervated motor units may remodel through reinnervation by nearby lower-threshold motor neurones [4,61], which may explain the reduced recruitment threshold observed in the older adults (as discussed below). These detrimental alterations in motor neurone structure are associated with reduced Ca^2+^-mediated plateau potential durations in striatal neurones (from aged rats) [62], slower conduction velocity of efferent motor axons [63], lower incidence of doublet discharges, slower maximum discharge rates [13,14,64] alongside an increased afterhyperpolarisation duration [16], and, as evidenced in the present study, lower PIC amplitude in the motor neurones. It is a logical hypothesis that the changes within the monoaminergic system and motor neurone structural integrity would possibly explain the reduced ΔFs observed in the present study. However, our findings indicate the need for further study of the cause-effect relationship between these mechanisms and reduced PICs in humans.

PICs are highly sensitive to synaptic inhibition, and both reciprocal and recurrent inhibition directly influence intrinsic motor neurone excitability, being effective PIC deactivators by opposing the facilitatory effects on PICs of descending brainstem neuromodulatory systems [65–69]. It would therefore be expected that the reduced PIC amplitude estimates observed in the older adults in the present study might result from an age-dependent increase in reciprocal and/or recurrent inhibition. However, older adults present reduced reciprocal inhibition from the common peroneal nerve onto *soleus* and from the tibial nerve onto the *tibialis anterior* [70]. Also, recurrent inhibition onto the *soleus* motor neurone pool is suspected to be relatively unaffected by ageing [71]. Therefore, a reasonable supposition is that the behaviour of spinal inhibitory/excitatory systems (i.e., reduced reciprocal inhibition and preserved recurrent inhibition) of older adults are unlikely to be responsible for the decreases in PIC amplitude.

The age-related reductions in PIC amplitudes indicated by our data may partly explain losses in motor function with ageing, and may thus have important clinical practice implications. Motor neurone PICs can amplify the synaptic input they receive, allowing motor neurones to discharge at higher rates, as shown in animal, computational modelling, and human studies [10,11,29,72]. This amplification system is an important feature when higher-intensity muscular contractions are needed. Thus, weaker motor neurone PICs could be an important limiting factor which to some extent explains the lower voluntary activation levels [73] and force production [2] observed in older adults. A lower capacity for producing high forces as a result of weaker PICs could therefore account for the increase in relative intensity and level of effort needed for older adults when performing daily activities [74]. This would indicate a reduced performance and increased fatigability when performing these activities [75]. Moreover, if these assumptions are true then the improvements in force and functional capacity following resistance training in older adults [76,77] could be partly mediated by increases in PIC amplitudes [24]. However, these causal hypotheses remain untested.

### Recruitment threshold

Motor units that were identified in the older adults in our study showed a lower recruitment threshold and were thus recruited at a lower torque level than in young adults. This may be a result of two different factors: 1) the decomposition algorithm may have biased the identification of motor units towards the lower-threshold ones. Older adults have a greater proportion of lower-threshold motor units as a result of motor unit remodelling subsequent to motor neuronal denervation, so motor units previously innervated by higher-threshold motor units become reinnervated by lower-threshold motor units [61]. Therefore, the decomposition may have picked up more low-threshold motor units in older adults, whereas the higher-threshold units in young may bias the decomposition the opposite direction. 2) The observed lower recruitment thresholds might reflect the recruitment of a greater relative number of smaller motor neurones during the task. Since motor unit discharge rate modulation in response to force changes is impaired in older adults, additional motor units must be recruited earlier in a triangular-shaped contraction to continually increase muscle force [78,79]. If this is the case, our data in slow isometric contractions indicate that aged motor neurones have a constrained capacity to amplify the excitatory synaptic input, consequently demanding an earlier recruitment of additional motor units to achieve the required motor output.

It is important to note that Ca^2+^ PIC channels can be activated below the action potential threshold (i.e., subthreshold PICs), strongly influencing motor unit recruitment [80]. The possibility exists that the population of lower recruitment threshold motor units identified by the decomposition algorithm in the older adults presents a higher motor neurone excitability (i.e. enhanced motor neurone recruitment for a given input) because of stronger subthreshold PIC activation. An increase in the proportion of subthreshold PICs would reduce the recruitment threshold while also decreasing ΔF values. Subthreshold PICs may be stronger in smaller motor neurones, resulting in a larger overall (sub-plus supra-threshold) PIC amplitude [31]. However, the ΔF method only estimates the suprathreshold contribution of the PICs to the discharge behaviour of motor units. Consequently, it is not possible to ascertain the behaviour of subthreshold PICs using the ΔF method. Our analytical approach accounted for any effect of recruitment threshold (and possibly motor neurone size) on ΔF, that may otherwise confound any motor-unit-population-related effect, by including recruitment threshold in our modelling of ΔF. Further, the reanalysis of ΔF using the composite paired motor unit method appeared to remove the effect of recruitment threshold on ΔF values (from *β* = 0.47, p < .001 to *β* = −0.05, p = .61), which is an expected characteristic of the method [31]. Therefore, the reduced ΔFs observed in older adults in the present study is not likely to have been an artefact of the different populations of motor units identified between groups.

### Peak discharge rates

The lower ΔF values in the older adults were also accompanied by reduced motor neurone discharge rates. The PIC is an important modulator of discharge rate output [8,29] and the available monoamines, serotonin and noradrenaline, facilitate PICs to increase motor neuronal gain and alter the input-output relationship according to the required output [25,29,81–84]. PICs can amplify synaptic input by more than five-fold, and are thus a determinant mechanism influencing the capacity for motor neurones to achieve the necessary discharge rates to obtain very high muscle activation levels [10–12]. We have recently shown that increases in discharge rate during triangular-shaped contractions at different force levels were strongly associated with ΔF increases, and thus PIC amplitudes, using a within-subject design [29]. Data from the current study reveal an important contribution of ΔF to motor neurone discharge rates, in which ΔF explained 53% of the variance in peak discharge rate. Motor unit discharge rates are not only modulated by PICs, but also depend on the ionotropic input they receive. Evidence in both animal [85,86] and human models [87] suggests that the synaptic input onto the motor neurone decreases with ageing, which may result from increases in intracortical inhibition or reduced intracortical facilitation in older adults [88–90]. Therefore, the lower peak discharge rates observed in older adults is a result of the lower PIC amplitudes and reduced synaptic input (descending drive and afferent feedback) received by the motor neurones.

### Strengths and limitations

The main strength of our study was the use of two validated [8,25,31] and a widely used methods to estimate PIC strength in humans [8,25,29,36,38,44,91]; however, both methods have limitations that should be pointed out [26]. The pairwise method [8] allowed us to obtain several pairs of motor units, having a larger amount of test units per participant. On the other hand, this method present a higher variance as a result of under and overestimation of ΔFs as a consequence of adopting control units with varied recruitment thresholds [31]. Recently, Afsharipour et al., [31] proposed the use of a composite control unit to reduce the ΔF variance present in the conventional paired motor unit analysis. However, this method requires some additional assumptions, such as the overlay of three lower threshold motor units with a similar discharge rate profile and with recruitment threshold below 3%. When following these assumptions, there was an important reduction of motor unit pairs, particularly for *soleus.* Therefore, we initially adopted the pairwise method as it permitted comparison between *soleus* and *tibialis anterior* ΔFs. We also ran the composite method as an additional analysis to examine whether reducing ΔF variability and removing the influence of the recruitment threshold on ΔF obtained with the pairwise method would affect our main outcomes. Using both methods allowed us to identify a large difference in ΔF between young and older adults, minimising the methodological limitations. Furthermore, it is worth mentioning that the assessors were not blinded to age group when visually inspecting and editing the motor units and future studies should confirm this with blinded assessors.

The lower recruitment threshold observed in the older adults may have been a result of bias of the decomposition algorithm because of the greater proportion of this type of motor units in this population. Therefore, caution should be taken when interpreting the differences in the recruitment threshold of motor units identified between these two heterogeneous groups. Indeed, our analytical approach included the variable recruitment threshold when modelling ΔF aiming to control any potential confounding factor related to potential different physiological behaviours of distinct populations of motor units.

Furthermore, ΔF values obtained in our study are derived from lower threshold *soleus* and *tibialis anterior* motor units recruited at a low force level (20% of peak torque); this is a commonly-used force target and is also similar to forces that might be expected in daily activities such as standing [39] and walking. However, older adults perform activities such as chair rise and both stair ascent and descent at a higher level of effort relative to their maximum capability [74] than young adults. Consequently, the ΔF data obtained during low force levels might not represent the motor neurone PIC behaviour at these daily activities requiring higher intensity contractions. Indeed, there is evidence that the function and structure of higher threshold motor neurones are more affected than lower threshold motor neurones [1]. This could hypothetically indicate a greater impairment in PICs during higher intensity contractions. Moreover, daily activities require the activation of different muscle groups. Motor neurones from distinct muscles depict different discharge behaviours during ageing [14]; thus, it is possible that ΔF behaviour might also differ. Therefore, we recommend that future studies investigate the effect of ageing on ΔF values from different muscles and at different contraction intensities, and its influence on possible impairments in physical function.

The cross-sectional design and the small group of tested individuals are additional limitations inherent to our study that should be mentioned. A cross-section study does not allow one to parse out causation *per se.* Longitudinal studies would involve repeated data collection from the same sample over several years to provide a better understanding of the effects of ageing on ΔF. In addition, the small group of non-sarcopaenic older adults with normal mobility tested in the present study does not allow our results to be extrapolated to populations with different characteristics or health conditions, such as very old (>85 years old), sarcopaenic, frail, or those with neurological disorders. They may also not represent individuals who consistently perform high levels of physical activity. Having a broader spectrum of older adults with low to high physical function and force levels would allow a more adequate investigation of the relationship between ΔF with these parameters.

## Conclusions

The present study provides novel evidence of reduced intrinsic motor neurone excitability in a group of non-sarcopaenic older adults with normal mobility by estimating PIC amplitudes using the paired-motor unit analyses. Older adults had substantially lower ΔFs, and presumably PIC amplitudes, in both *soleus* and *tibialis anterior* than young adults with comparable physical activity level. This would likely influence the capacity of older individuals to activate the muscles, thus requiring a greater descending drive from cortical areas and hence level of volitional effort, and greater number of recruited motor units to achieve the same force level (relative to maximum). We also identified a small contribution of ΔF to the between-subject variability in peak discharge rates. The present findings contribute to our understanding of the effects of ageing on motor neurone excitability, which is a potential mechanism underpinning motor functional loss during ageing; this hypothesis should be explicitly tested in future studies. Two logical next steps are: 1) to examine the effect of ageing on monoaminergic projections onto the motor neurones and their relationship with the reduced PIC amplitude observed in the present study; 2) to investigate the association between ΔF values for motor units in different muscles and the variance in performance on clinical tests of motor function.

## ADDITIONAL INFORMATION

### Data and code availability

The dataset and R code are available at https://github.com/orssatto/PICs-ageing.

### Author contributions

LBRO, AJS, AJB, and GST contributed with the conception and design of the work. LBRO acquired data. RLS developed the MATLAB script for ΔF, peak discharge rate, and recruitment threshold calculation. LBRO conducted the biological signals data analyses, and DNB developed the R script and conducted statistical analyses. All authors interpreted and discussed the data, drafted the manuscript, and revised it critically providing important intellectual content.

All authors approved the final version of the manuscript; agree to be accountable for all aspects of the work in ensuring that questions related to the accuracy or integrity of any part of the work are appropriately investigated and resolved; and qualify for authorship, and all those who qualify for authorship are listed.

