## Supplemental material 1 for "INTRINSIC MOTOR NEURONE EXCITABILITY IS REDUCED IN SOLEUS AND TIBIALIS ANTERIOR OF OLDER ADULTS"

**Supplement 1.** Candidate models considered for the analysis of ΔF.

| Model Description | Model set up | BIC value |
| --- | --- | --- |
| Recruitment threshold only | ΔF ~ age_group + muscle + recruit_s + age_group*muscle + (1 + recruit_s \| participant) | 2069.9 |
| Recruitment threshold by muscle interaction | ΔF ~ age_group + muscle + recruit_s + age_group*muscle + muscle*recruit_s + (1 + recruit_s*muscle \| participant) | 2060.8 |
| Recruitment threshold by age interaction | ΔF ~ age_group + muscle + recruit_s + age_group*muscle + age_group*recruit_s + (1 + recruit_s \| participant) | 2075.9 |
| All 2-way interactions | ΔF ~ age_group + muscle + recruit_s + age_group*muscle + age_group*recruit_s + muscle*recruit_s + (1 + recruit_s*muscle \| participant) | 2065.5 |
| All 2-way interactions and the 3-way interactions | ΔF ~ age_group + muscle + recruit_s + age_group*muscle + age_group*recruit_s + muscle*recruit_s + age_group*muscle*recruit_s + (1 + recruit_s*muscle \| participant) | 2070.3 |

*Note*. BIC = Bayesian Information Criteria, recruit_s = Recruitment threshold (standardised; mean = 0, standard deviation = 1).

Smaller Bayesian Information Criteria values indicate a better fitting model.
